# Catestatin improves insulin sensitivity in diet-induced obese mice: *in vivo* and *in silico* validation

**DOI:** 10.1101/615997

**Authors:** Abhijit Dasgupta, Keya Bandyopadhyay, Indrani Ray, Gautam K. Bandyopadhyay, Nirmalya Chowdhury, Rajat K. De, Sushil K. Mahata

## Abstract

Obesity is characterized by a state of chronic, unresolved inflammation in insulin-targeted tissues. Obesity-induced inflammation accumulates proinflammatory macrophages in adipose tissue and liver. Proinflammatory cytokines released from tissue macrophages inhibits insulin sensitivity. Chromogranin A (CgA) peptide catestatin (CST: hCgA_352-372_) improves obesity-induced hepatic insulin resistance by reducing inflammation and inhibiting proinflammatory macrophage infiltration. Obesity leads to inflammation-induced endoplasmic reticulum (ER) stress and insulin resistance. We reasoned that the anti-inflammatory effects of CST would alleviate ER stress. CST decreased obesity-induced ER dilation in hepatocytes and macrophages. CST reduced phosphorylation of UPR signaling molecules and increased phosphorylation of insulin signaling molecules. We developed an *in silico* state space model mimicking dynamics of integrated ER stress and insulin pathways. Proportional-Integral-Derivative (PID) controllers helped in checking whether the reduction of phosphorylated PERK resulting in attenuation of ER stress, resembling CST effect, could enhance insulin sensitivity. The simulation results showed CST not only decreased ER stress but also enhanced insulin sensitivity. Simulation results also revealed that enhancement of AKT phosphorylation overcame effects of high ER stress to achieve insulin sensitivity.

## Introduction

The liver maintains whole body homeostasis by regulating critical metabolic, secretory and excretory functions. Calcium storage, protein and lipid synthesis along with protein folding are the key functions of the endoplasmic reticulum (ER) (*1*). Hepatocytes (representing up to 70% of total liver cells) contain both rough and smooth ER, which perform the myriad of metabolic functions (*2*). The smooth ER synthesizes not only the majority of the membrane lipids but also their intermediates such as cholesterol, ceramides, and glycerophospholipids (*3*, *4*). Both rodents (*5*, *6*) and humans (*7*–*9*) accumulate ceramides in tissues and plasma (*10*), which inhibits insulin action by decreasing phosphorylation of AKT and consequent inhibition of glucose uptake. Ceramide also activates nuclear factor-κ-B (NF-κB)-tumor necrosis factor-α (TNF-α) axis and induces inflammation (*5*, *6*). The rough ER controls the synthesis and maturation of proteins, which comprises up to 40% of cells proteome of the secretory pathway (*11*). Ribosomes perform the translation of proteins on the cytosolic surface of the ER (*12*), and sec61 complex translocates the unfolded polypeptide into the ER lumen (*13*) where they undergo N-glycosylation and folding into secondary or tertiary structures. The rough ER lumen is enriched with high concentrations of calcium, molecular chaperones and folding enzymes, which facilitates protein folding and maturation (*14*). Non-native proteins are recognized by the ER associated degradation (ERAD) quality control system and are degraded by the cytosolic ubiquitin-proteasome system (*15*, *16*). ER stress is characterized by the accumulation of misfolded or unfolded proteins in response to environmental insults, increased protein synthesis and reduced secretory efficacy (*17*, *18*). Homeostasis is restored by the ER stress-induced activation of the adaptive unfolded protein response (UPR). The following three ER localized proteins initiate UPR signalling in mammalian cells: double-stranded RNA-dependent protein kinase-like ER kinase (PERK) – eukaryotic translation initiation factor 2α (eIF2α), inositol-requiring 1α (IRE1α) – X-box-binding protein (XBP1), and activating transcription factor-6α (ATF6α) (*19*). When physiological UPR becomes chronically activated, ER stress occurs. Thus, the chronic activation of the UPR has been reported in human obesity and non-alcoholic fatty liver disease (NAFLD), and in the adipose and/or liver tissue of dietary and genetic murine models of obesity (*20*–*26*).

The levels of free fatty acids (FFA), insulin, glucose, proinflammatory cytokines and ceramides are increased in blood of obese rodents and humans, which activates the innate immune system resulting in a chronic low-grade inflammation of white adipose tissue (*10*, *27*) and the subsequent development of insulin resistance on other peripheral tissues, including the skeletal muscle, adipose and liver (*28*–*30*). Thus, obesity aggravates both inflammation and ER stress.

A recent investigation has shown that the chromogranin A (CgA) peptide catestatin (CST: human CgA_352-372_) improves hepatic insulin sensitivity in diet-induced obese (DIO) mice as well as in insulin-resistant CST knockout mice by reducing inflammation and inhibiting infiltration of macrophages (*31*). Since ER stress activates the inflammatory response (*32*–*34*) and the inflammatory response in turn also activates ER stress (*35*), we reasoned that one additional mechanism by which CST can improve insulin resistance in DIO mice is by alleviating ER stress. We have tested the above hypothesis by looking at the UPR signaling in DIO liver as well as by developing an *in silico* state space model for mimicking similar cellular dynamics as shown by different experimental results under ER stress (DIO) and no ER stress (normal chow diet (NCD)). Besides, two Proportional-Integral-Derivative (PID) controllers were applied to control the signal level of tyrosine phosphorylated insulin receptor (IRpY), tyrosine phosphorylated insulin receptor substrate (IRSpY), and phosphorylated Protein Kinase B (pAKT) along with phosphorylated PERK (pPERK) to low/high values. The simulation results clearly indicated that reduction of pPERK resulting in attenuation of ER stress, which was achieved by applying CST as shown by the experimental results, led to high insulin sensitivity. Besides, higher level of IRpY and IRSpY were unable to enhance insulin sensitivity during high ER stress. Thus, the reduction of ER stress on application of CST is one of the potential factors to enhance insulin sensitivity in mammalian cells. However, our simulation predicts that enhancement of pAKT would enhance insulin sensitivity in spite of high ER stress.

## Materials and Methods

### Animals, Diets and Treatments

Diet-induced obese mice was created by feeding male wild-type (WT) mice (starting at 8 weeks of age) with a high-fat diet (HFD, Research diets D12492, 60% of calories from fat) for 16 weeks. Mice were kept in a 12:12 hour dark/light cycle; food and water was available at all times. Control mice were fed a NCD (14% of calories from fat). Mice were treated with CST (5 µg/g BW IP for 15 days) after 11 weeks of HFD feeding when weight gains practically levelled off. In accordance with NIH animal care guidelines, all procedures and animals were housed and handled with the approval of The Institutional Animal Care and Utilization Committee.

### Transmission Electron Microscopy (TEM)

WT-NCD, WT-DIO and WT-DIO+CST livers were perfusion fixed through the left ventricle under deep anaesthesia. A pre-warmed (37°C) calcium and magnesium free buffer consisting of DPBS (Life Technologies Inc. Carlsbad, CA), 10 mM HEPES, 0.2 mM EGTA, 0.2% bovine serum albumin, 5 mM glucose and 10 mM KCl was used to flush mice for 3 min (3 ml per min; Langer Instruments Corp, Boonton, NJ). This is followed by perfusion with freshly prepared pre-warmed (37°C) fixative containing 2.5% glutaraldehyde, 2% paraformaldehydein 0.15 M cacodylate buffer for 3 min. After dissecting, liver slices (2 mm thick) were put in the same fixative overnight (2 hours at room temperature and 12 hours at 4°C), and postfixed in 1% OsO_4_ in 0.1 M cacodylate buffer for 1 hour on ice. Liver slices were stained *en bloc* with 2-3% uranyl acetate for 1 hour on ice followed by dehydration in graded series of ethanol (20-100%) on ice, one wash with 100% ethanol and two washes with acetone (15 min each) and embedded with Durcupan. Approximately, 50 to 60 nm thick sections were cut on a Leica UCT ultramicrotome. Sections were picked upon Formvar and carbon-coated copper grids and stained with 2% uranyl acetate for 5 minutes and Sato’s lead stain for 1 minute. Livers (3 from each group) were fixed and processed in two separate days. Stained grids were looked under a JEOL 1200EXII (JEOL, Peabody, MA) TEM and photographed with a Gatan digital camera (Gatan, Pleasanton, CA). Random micrographs were taken from 3 livers where samples were blinded. Also, 2 people did measurements randomly from different tissues as described previously (*36*). The width of the ER lumen was determined by using the free-hand tool in NIH ImageJ 1.49 software.

### Immunoblotting

Homogenization of livers was made in a lysis buffer supplemented with phosphatase and protease inhibitors. Homogenates were subjected to 10% SDS-PAGE and immunoblotted. The following primary antibodies were obtained from Cell Signaling Technology (Boston, MA): IRE (rabbit polyclonal 1:1000) and pS724-IRE (rabbit polyclonal 1:500), PERK (rabbit polyclonal 1:1000) and pT980-PERK (rabbit polyclonal 1:1000), eIF (rabbit polyclonal 1:1000) and pS51-eIF (rabbit polyclonal 1:500).

### Real Time PCR

RNeasy Mini Kit (Qiagen) was used to isolate total RNAfrom liver tissues. qScriptcDNA synthesis kit (QuantaBio, Beverly, MA) was used to make the cDNAs, which were amplified by using PERFECTA SYBR FASTMIX L-ROX1250 (QuantaBio). The amplified samples were run on an Applied Biosystems 7500 Fast Real-Time PCR system (ABI). All PCRs were normalized to *Gapdh*, and relative expression levels were determined by the ΔΔ*C_t_* method. Primer sequences are provided in Table S1 of Supplementary File S1.

### Computational analyses

The computational model involved formulation of state transition equations, input design, estimation of values for the kinetic parameters through model validation, and finally, application of PID controllers. In order to formulate state transition equations, we integrated ER stress and insulin signaling pathway as depicted in Fig. S1 of Supplementary File S1 using the method in one of our previous investigations (*37*). Using these state transition equations together with estimated input and parameter values, the *in silico* state space model was developed. In order to investigate the CST effects on insulin sensitivity along with alleviation of ER stress, we applied PID controllers on the state space model as depicted in Fig. S2 of Supplementary File S1.

### Formulation of state transition equations

Let us assume that the integrated biochemical pathway (Fig. S1 of Supplementary File S1) under investigation involves the state components *x*_1_, *x*_2_,…, *x*_n_ representing different signaling molecules/proteins. Here, *u*_1_ and *u*_2_ are external inputs representing ER stress and insulin. Let *x*_1_ is triggered by *u*_1_. Here, *x*_1_ is decayed/consumed at the rate proportional to *x*_1_×*x*_2_. Thus, we can write,

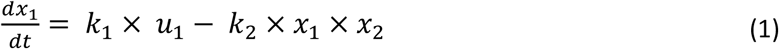

Here *k*_1_ and *k*_2_ are interaction rate and decay constants respectively.

Let us consider a situation as an example, where *x*_20_ is activated by *x*_19_ under the influence of *x*_1_ and *x*_2_. This activation is accelerated by *x*_18_. Besides, *x*_20_ is decayed/consumed at the rate proportional to *x*_20_. Here, we can write,

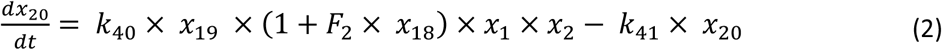

Here *k*_40_ and *k*_41_ are interaction rate and decay constants respectively, whereas *F*_2_ is binding constant. Again, *x*_26_ is triggered by *u*_2_, and decayed/consumed at the rate proportional to *x*_26_ along with other feedback effects due to *x*_30_ and *x*_32_. Thus, we can write,

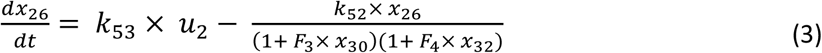

Here *k*_53_ and *k*_52_ are interaction rate and decay constants respectively, whereas *F*_3_ and *F*_4_ are binding constants. Similarly, we developed state transition equations for all other state components according to the biological phenomena. We have included all state transition equations in Supplementary File S1 to restrict the length of manuscript.

### Inputs u_1_ and u_2_

In order to get the equations for external inputs, we considered system equilibrium (steady state) condition. At this condition, we can say 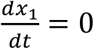 and 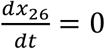. Thus, we have

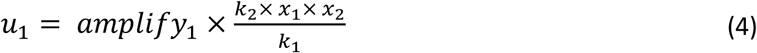

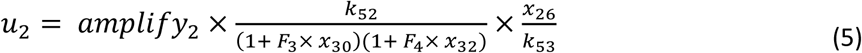

Here, *amplify*_1_ and *amplify*_2_ are constant terms. They are used to amplify the effect of external inputs to the system.

### Estimation of the values of the constant parameters through model validation

We solved the ordinary differential equations (ODEs), formulated above, with random values for the constant parameters (interaction rate, decay and binding constants) in [0, 5]. We initialized all state components at 1.05 and restrict their values in [1, 5] as depicted in Supplementary File S1. This ODE system was solved numerically (70000 iterations and in time span of [0 0.0002]) using ode23tb solver of Matlab software. As a result, the computational model was able to capture the behavioral pattern of different molecules (state components) under consideration at continuous time points. This is the advantage of ODE based model which is capable of calculating values of state components at continuous time points depending on the initial values provided in the beginning of simulation. The computational results were validated to check whether the model follows the experimental behavioral patterns or not for both DIO and NCD situations. Although the experimental western blot results provided molecular expression/concentration level (low/high) at single time point, they were quite capable of providing information about the change of molecular expression/concentration level during DIO and NCD situations. Based on this experimental knowledgebase, if the result did not follow the experimental patterns, we changed the values of the constant terms in an ad hoc manner to replicate the experimental behavior. Whenever the model followed the experimental behavioral patterns, we fixed the values of the constant parameters as provided in Supplementary File S1. Here we considered two situations-one calculating the value of *u*_1_ (ER stress) using Equation (4) resembling stress (DIO) and the other depicting *u*_1_ (ER stress) = 0 as control (NCD).

### Application of PID controllers to investigate CST effect

Here we applied two PID controllers on the ODE based model (state space model) considering it as a plant to test if the attenuation of ER stress resembling experimental CST effect and the enhancement of insulin sensitivity can be achieved simultaneously. In order to accomplish that, the PID controllers were involved in controlling the signal levels of pPERK, IRpY, IRSpY and pAKT. Here we used Simulink platform of Matlab software for simulation. Depending on error functions, appropriately calculated control (external) inputs for ER stress and insulin were applied on the state space model. The general form of the error function can be defined as

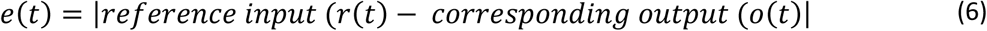

Finally we considered the tuned parameter values as *k*_p_ = 0, *k*_i_ = 0.0691 and *k*_d_ = 0 for first PID controller. While for the second PID controller, we tuned parameter values as *k*_p_ = 0.2134, *k*_i_= 0.10329 and *k*_d_ = −0.1082. Here *k*_p_, *k*_i_ and *k*_d_ represent proportional, integral and derivative constants respectively. The general form of the control (external) input function (*u(t)*) can be defined as

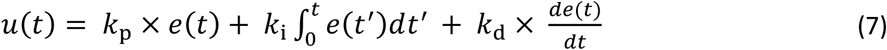

## Results

### Hepatocyte ER dilation and activation of the ATF6 branch of the UPR pathway

The increased demand on the synthetic machinery during obesity results in unfolded or misfolded proteins accumulation in the ER lumen leading to ER stress, activating the UPR. Our ultrastructural studies show dilated ER lumen possibly to accommodate increased unfolded/misfolded proteins (Fig. 1A-C). ATF6α is an ER type-II transmembrane protein harbouring a bZIP transcription factor on its cytosolic domain and a C-terminal luminal region that senses ER stress (Fig. 1D). To counteract with the ER stress, ATF6α transits to the Golgi apparatus for its cleavage by the proteases S1P and S2P, which releases the cytosolic domain (ATF6f) (*38*). ER’s capacity for folding is increased as a result of ATF6f-induced expression of protein chaperone genes including ER protein 57 (ERp57), binding immunoglobulin protein (BiP) and glucose-regulated protein (GRP) 74. ATF6f also decreases lipogenesis in liver by antagonizing SREBP2 (*39*) and inhibits gluconeogenesis by interacting with CRTC2 (*40*) or inhibition of CREB (*41*). Consistent with the existing literature, we found increased ATF6α mRNA level in DIO liver (*42*, *43*) (Fig. 1E).

**Figure 1:**
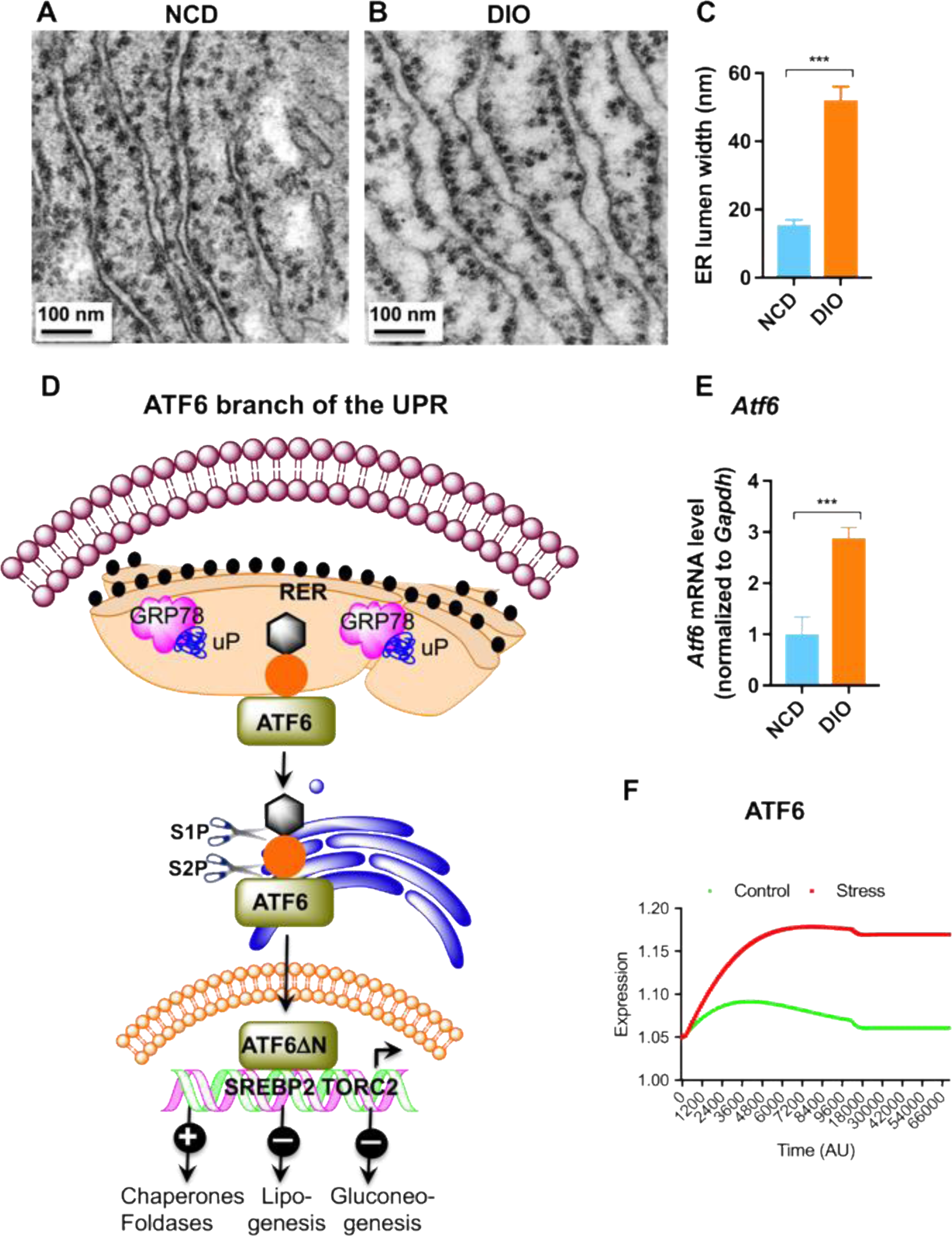
(A&B) TEM photographs showing dilation of ER lumen in ultrathin (~60 nm) liver sections of DIO mice. (C) Morphometric analyses of ER lumen width in NCD and DIO mice. (D) Schematic diagram showing the ATF branch of the UPR. (E) Changes in ATF6α mRNA levels in NCD and DIO mice (n=6). (F) *In silico* state space model resembling the behavior of ATF6α with the experimental results. The expression of ATF6α is higher during stress (DIO) condition than that in control (NCD). **p<0.001.

Depending on the aforesaid experimental results, we validated the *in silico* state space model during stress (DIO) and control (NCD) situations. We found higher expression of ATF6α during stress (DIO) than that in control (NCD) after 70000 iterations (Fig. 1F), which corresponds to increased ATF6α mRNA in DIO liver (Fig. 1E).

### Activation of the IRE1α branch of the UPR

The transmembrane protein IRE1α contains an N-terminal luminal sensor domain, a C-terminal cytosolic effector, and a single transmembrane domain harbouring a Ser/Thrkinase domain and an endoribonuclease domain (Fig. 2A). ER stress induces dimerization and trans-autophosphorylation of IRE1α, thereby causing an induction of a conformational change and the subsequent activation of its RNAse domain to selectively cleaves a 26-nucleotide intron within the *XBP1* mRNA (*44*, *45*). This unconventional splicing introduces a translational frameshift to generate a stable and active transcription factor XBP1s. Active XBP1s regulate the expression of genes that modulate protein folding, secretion, translocation, ERAD into the ER, and synthesis of lipid (*46*, *47*). To alleviate the protein-folding load on the ER, the RNAse domain of IRE1α also modulates the “regulated IRE1-dependent decay (RIDD)” (*48*). IRE1α also induces lipogenesis (*49*, *50*) and gluconeogenesis (*51*). Consistent with the existing literature, we found increased phosphorylation of IRE1α (pIRE1α) in DIO liver (Fig. 2B). The state space model also showed higher signal of pIRE1α during stress (DIO) than that in control (NCD) (Fig. 2C), which corresponds to increased phosphorylated IRE1α in DIO liver (Fig. 2B).

**Figure 2:**
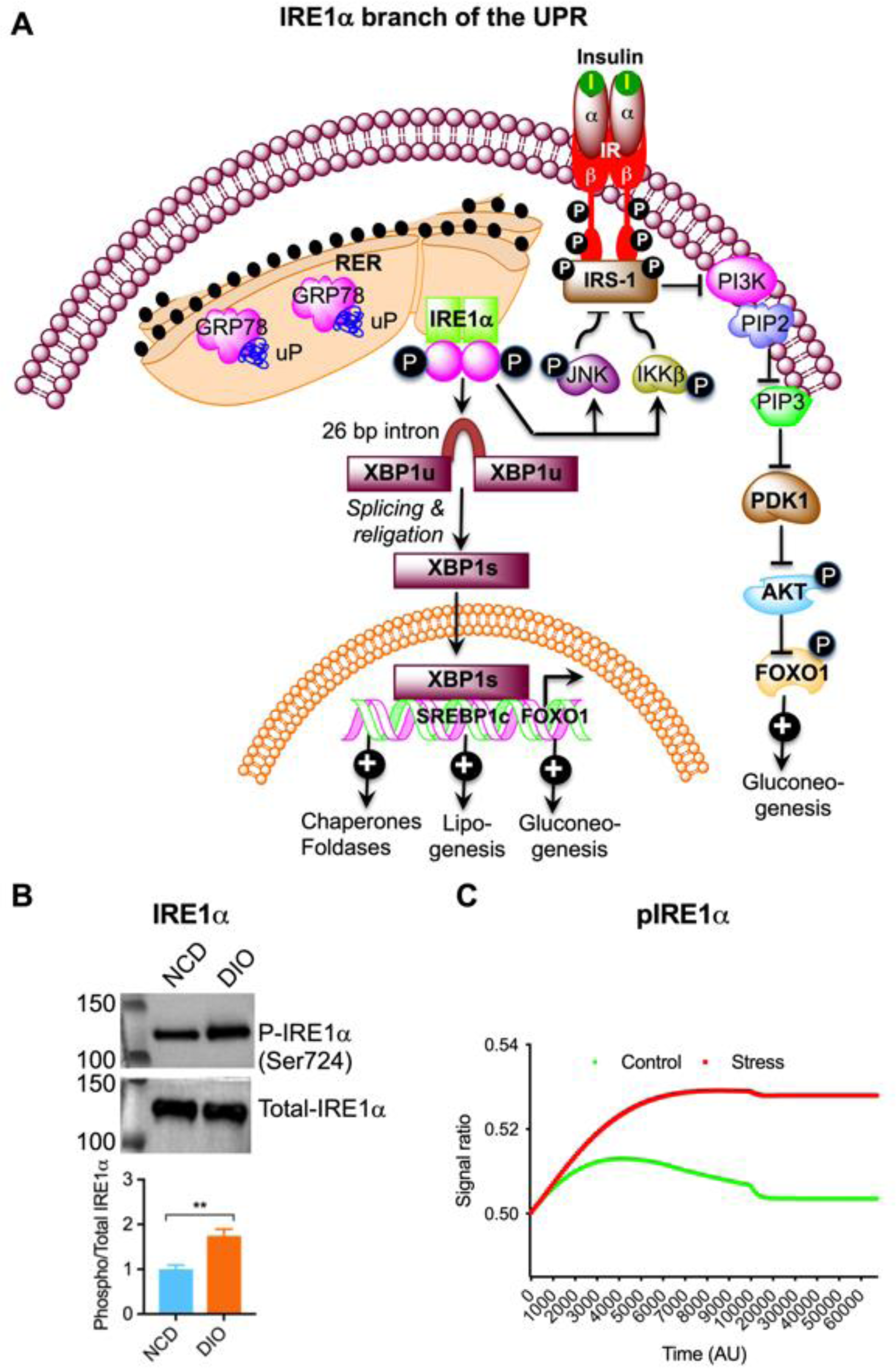
(A) Schematic diagram showing the IREα branch of the UPR. (B) Western blots showing increased phosphorylation of IRE1α in DIO liver (n=4). (C) *In silico* state space model resembling the behavior of pIRE1α with the experimental results. The ratio (pIRE1α/IRE1α) is higher during stress (DIO) condition than that in control (NCD). **p<0.01.

### Activation of the PERK branch of the UPR

Phosphorylation at Thr980 activates PERK to alleviate the protein-folding load on the ER. Phosphorylated PERK phosphorylates eIF2α at Ser51 to briefly halt the initiation of mRNA translation. This leads to the reduction of global protein synthesis resulting in decreased workload of the ER (Fig. 3A) (*52*, *53*). Paradoxically, phosphorylated eIF2α (peIF2α) also up regulates the transcription and translation of many mRNAs, such as nuclear erythroid 2 p45-related factor 2 (Nrf2), activating transcription factor-4 (ATF4), and nuclear factor kappa b (NF-κB). ATF4, produced through alternative translation, influences gene expressions involved in ER redox control (ERO1, ER oxidoreductin), apoptosis (CHOP, C/EBP homologous protein), glucose metabolism (fructose 1,6-bisphosphate; glucokinase, and phosphoenolpyruvate carboxykinase), and the negative feedback release of eIF2α inhibition (Gadd34, growth arrest, and DNA damage-inducible protein) (Fig. 3A) (*54*, *55*). Consistent with the above literature, we found increased phosphorylation of PERK at Thr980 (Fig. 3B) and eIF2α at Ser51 (Fig. 3C) as well as increased mRNA level of ATF4 (Fig. 3D).

**Figure 3:**
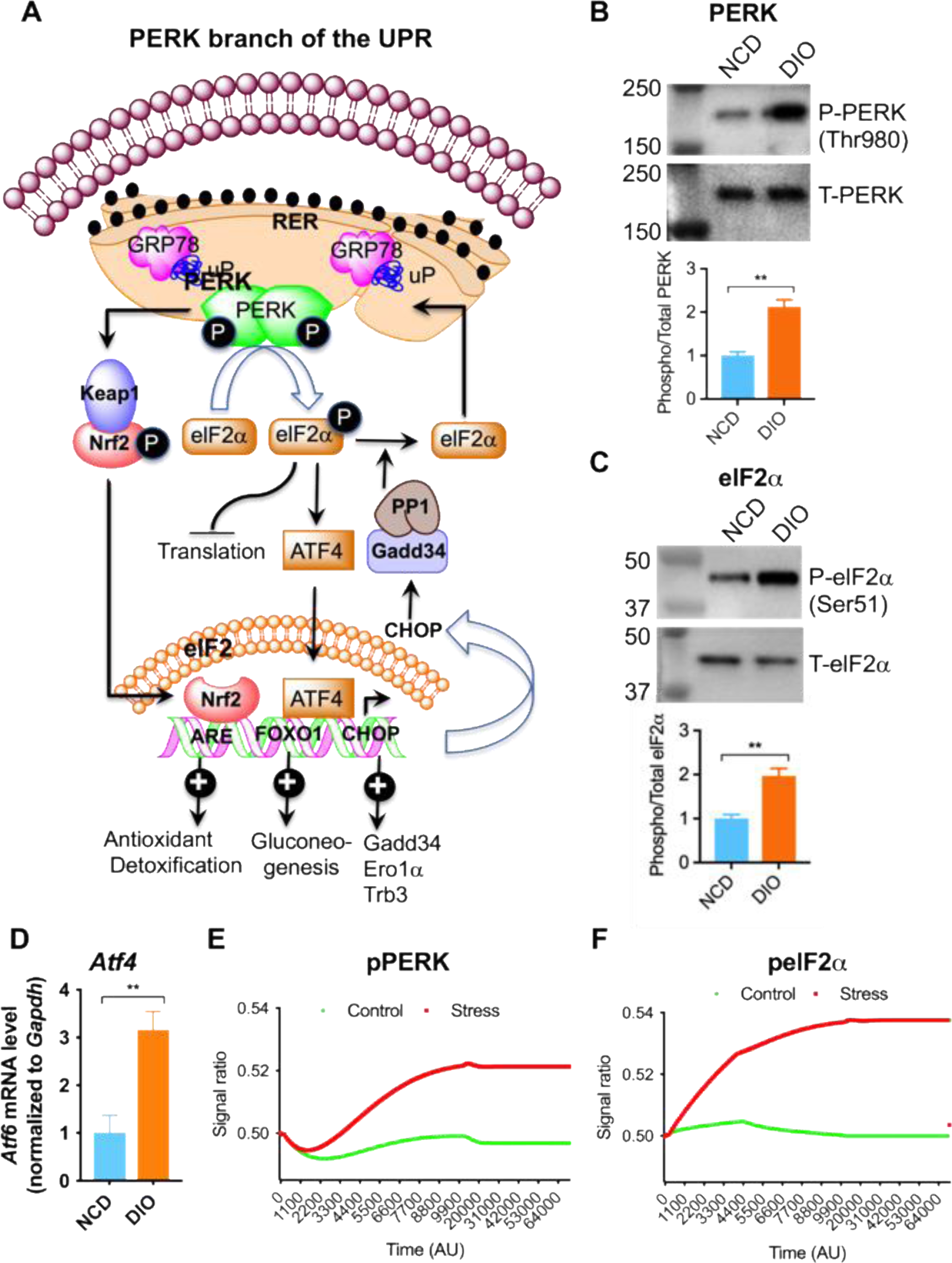
(A) Schematic diagram showing the PERK branch of the UPR. (B) Western blots showing increased phosphorylation of PERK at Thr980 in DIO liver (n=4). (C) Western blots showing increased phosphorylation of elF2α at Ser51 in DIO liver (n=4). (D) Changes in ATF4 mRNA level in NCD and DIO liver (n=6). (E&F) *In silico* state space model resembling the behavior of pPERK and peIF2α with the experimental results. The ratios (pPERK/PERK) and (pelF2α/pelF2α) are higher during stress (DIO) condition than that in control (NCD). **p<0.01.

The *in silico* state space model was quite effective to mimic the aforesaid experimental results. The model showed higher signals of pPERK (Fig. 3E) and peIF2α (Fig. 3F) during stress (DIO) than that in control (NCD), which corresponds to increased phosphorylated PERK (Fig. 3B) and phosphorylated eIF2α (Fig. 3C) in DIO liver.

### CST decreases ER stress

In lean state, adipose tissue macrophages (ATM) and resident macrophages (Kupffer cells) in liver exhibit an anti-inflammatory alternatively activated (M2) phenotype (*56*–*60*). Complex molecular interactions between diet, environment and genetics at the metabolic tissues (adipocytes, heaptocytes, and pancreatic islets) and immune system (macrophages, neutrophils, and lymphocytes) provoke a low-grade, chronic inflammatory response, which is called metaflammation (*61*). Obesity, characterized by metaflammation (*62*–*64*), is allied with ER stress that disrupts glucose homeostasis (*2*, *20*, *65*) and results in the development of atherosclerotic plaques (*66*–*68*). In obesity, ATMs and Kupffer cells display a predominantly proinflammatory classically activated (M1) phenotype, which is thought to promote insulin resistance and type 2 diabetes (*56*, *69*–*74*). It has been recently shown that IRE1α mediates saturated fatty acid-induced activation of the NLRp3 inflammasome in human and mouse macrophages (*75*) and that macrophage-specific deletion of IRE1α conferred resistance to high-fat diet-induced obesity, thereby linking macrophages to ER stress, metaflammation and insulin sensitivity (*76*). We have recently shown that CST improves insulin sensitivity by inhibiting obesity-induced inflammation and macrophage infiltration in the liver and by suppressing glucose production in hepatocyte (*31*). Therefore, we reasoned that CST would decrease obesity-induced ER stress. Consistent with our hypothesis we found that CST treatment exhibits ER stress lowering effects: (i) decrease of obesity-induced ER dilation (Fig. 4A-C), (ii) decreased mRNA abundance for ATF6 (Fig. 4D) and ATF4 genes (Fig. 4E), (iii) decreased abundance of spliced *Xbp1* mRNA (Fig. 4F), and (iv) decreased phosphorylation of PERK (Fig. 4G), eIF2α (Fig. 4H), and IRE1α (Fig. 4I).

**Figure 4:**
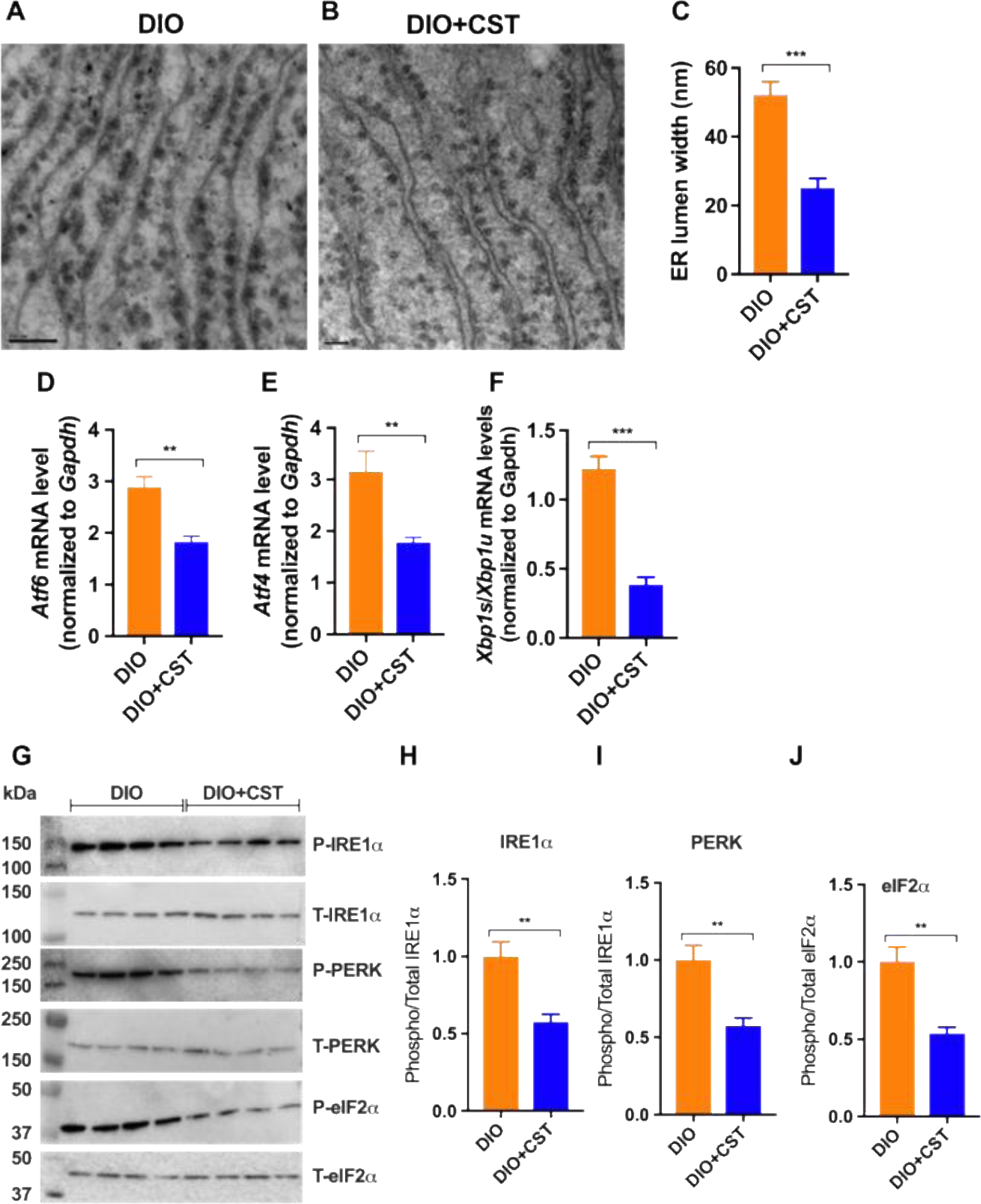
(A&B) TEM photographs showing attenuation of ER lumen in DIO liver sections after treatment with CST. (C) Morphometric analyses of TEM photographs showing decreased ER lumen width after treatment with CST. qPCR analyses showing CST-induced decrease in mRNA levels of (D) ATF6α, (E) ATF4, and (F) ratio between spliced versus unspliced *Xbp1* in DIO liver. Western blots showing decreased phosphorylation of UPR signaling molecules in DIO liver after treatment with CST: (G) phosphorylated PERK/total PERK, (H) phosphorylated elF2α/total elF2α, and (I) phosphorylated IRE1α/total IRE1α.

### CST decreases ER dilation in macrophages

Since macrophage-specific deletion of IRE1α conferred resistance to high-fat diet-induced obesity, thereby linking macrophages to ER stress, metaflammation and insulin sensitivity (*76*), we looked at the ER and found that the ER lumen got dilated (Fig. 5A-C & G), which was reduced markedly by CST (Fig. 5D-G).

**Figure 5:**
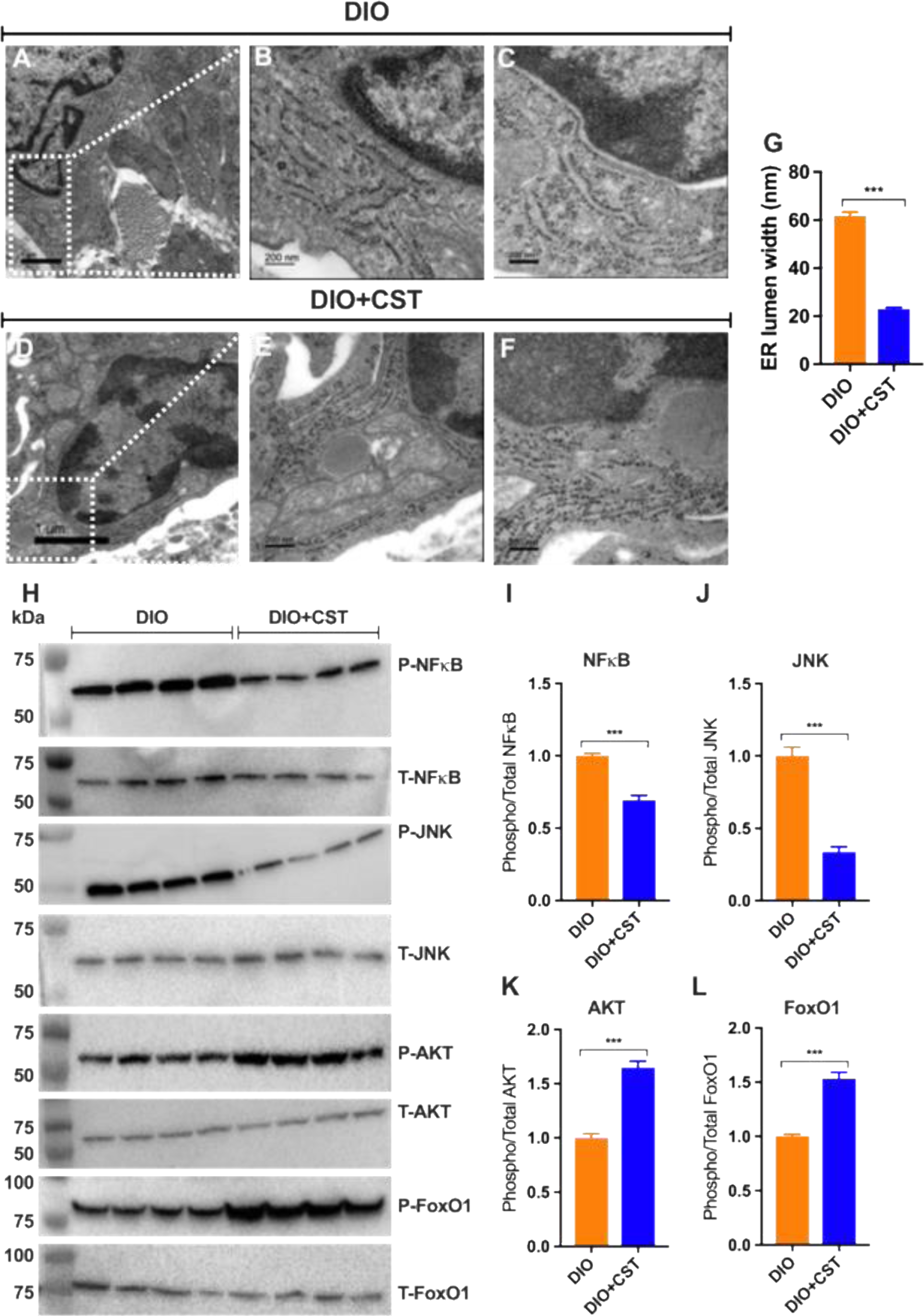
TEM photographs showing ER in infiltrated macrophages in DIO liver sections after treatment with CST. (A) Low magnification and (B&C) high magnification micrographs showing ER dilation in DIO mice. (D) Low magnification and (E&F) high magnification micrographs showing ER dilation in DIO mice treated with CST. (G) Morphometric analyses of ER lumen diameter in DIO and DIO+CST-treated mice. Western blots showing (H) decreased phosphorylation of NF-κB and (H) JNK coupled with (H) increased phosphorylation of AKT and FoxO1 in DIO liver after treatment with CST. The corresponding ratios of densitometric values are shown as follows: (I) NF-κB, (J) JNK, (K) AKT, and (L) FoxO1.

### Alleviation of ER stress by CST results in attenuation of inflammation and improvement in insulin signaling

The transcription factor NF-κB is normally suppressed by its inhibitor, IκBα. When IKK (IκB kinase) is activated, it phosphorylates IκBα, which leads to its degradation resulting in activation of NF-κB (*77*, *78*). Activated NF-κB translocates to the nucleus and augments the expression of proinflammatory genes(*79*–*81*). ER stress-induced IRE1 physically interacts with IKK, leading to an increase in phosphorylation of IκBα and the concomitant decrease in total levels of IκBα, which results in activation of NF-κB (*82*). It appears from the existing literature that during ER stress basal IKK activity, retained by IRE1, and PERK-mediated translation inhibition act in concert to activate NF-κB. Chronic treatment of DIO mice with CST resulted in significant decrease in phosphorylation of NF-κB, indicating attenuation of inflammation (Fig. 5H & I). *While* tyrosine phosphorylation activates, serine phosphorylation of insulin receptor substrates (IRS) at specific serine residues inhibits insulin signaling. ER stress induces insulin receptor signaling through increasing the serine phosphorylation and decreasing the tyrosine phosphorylation of IRS-1 (IRSpY), leading to insulin resistance (*83*). One of the mechanisms by which ER stress could impair insulin action is by the activation of JNK through double-stranded RNA-dependent protein kinase (PKR) (*63*), via the IRE1/TRAF2/ASK1 (*84*) or the PERK/ERO1L/CHOP/IP_3_R/ASK1(*85*) pathways, all of which have been reported to impair insulin receptor signaling (*20*, *79*, *82*, *86*). We found significant decrease in phosphorylation of JNK after chronic treatment with CST (Fig. 5H&J), which indicates improvement in insulin signaling. The improvement in insulin signaling by CST is further strengthened by increased phosphorylation of AKT (Fig. 5H & K) and FoxO1 (Fig. 5H & L).

### Computational model showing stress (HFD)-changes in insulin and inflammatory signaling

Previous investigations (*84*, *87*) showed that ER stress could lead to insulin resistance through different ways including activation of pJNK and pIκB kinase β (pIKKβ). Consistent with the existing literature, the *in silico* state space model yielded the following: enhanced insulin sensitivity in control (NCD) compared to that under stress (DIO) (Fig. 6A); (i) increased levels of IRpY (Fig. 6B), and (ii) IRSpY (Fig. 6C), (iii) pAKT (Fig. S3A of Supplementary File S1), (iv) phosphorylated Forkhead box protein O1 (pFoxO1) (Fig. S3B of Supplementary File S1) and (v) increased concentration of phosphatidylinositol trisphosphate (PIP_3_) (Fig. 6D) in control (NCD) compared to that under stress (DIO) condition. On the other hand, the signal levels of pJNK (Fig. 6E), pIKKβ (Fig. 6F) and pNF-κB (Fig. S3C of Supplementary File S1) were higher during stress (DIO) than that in control (NCD) scenario.

**Figure 6:**
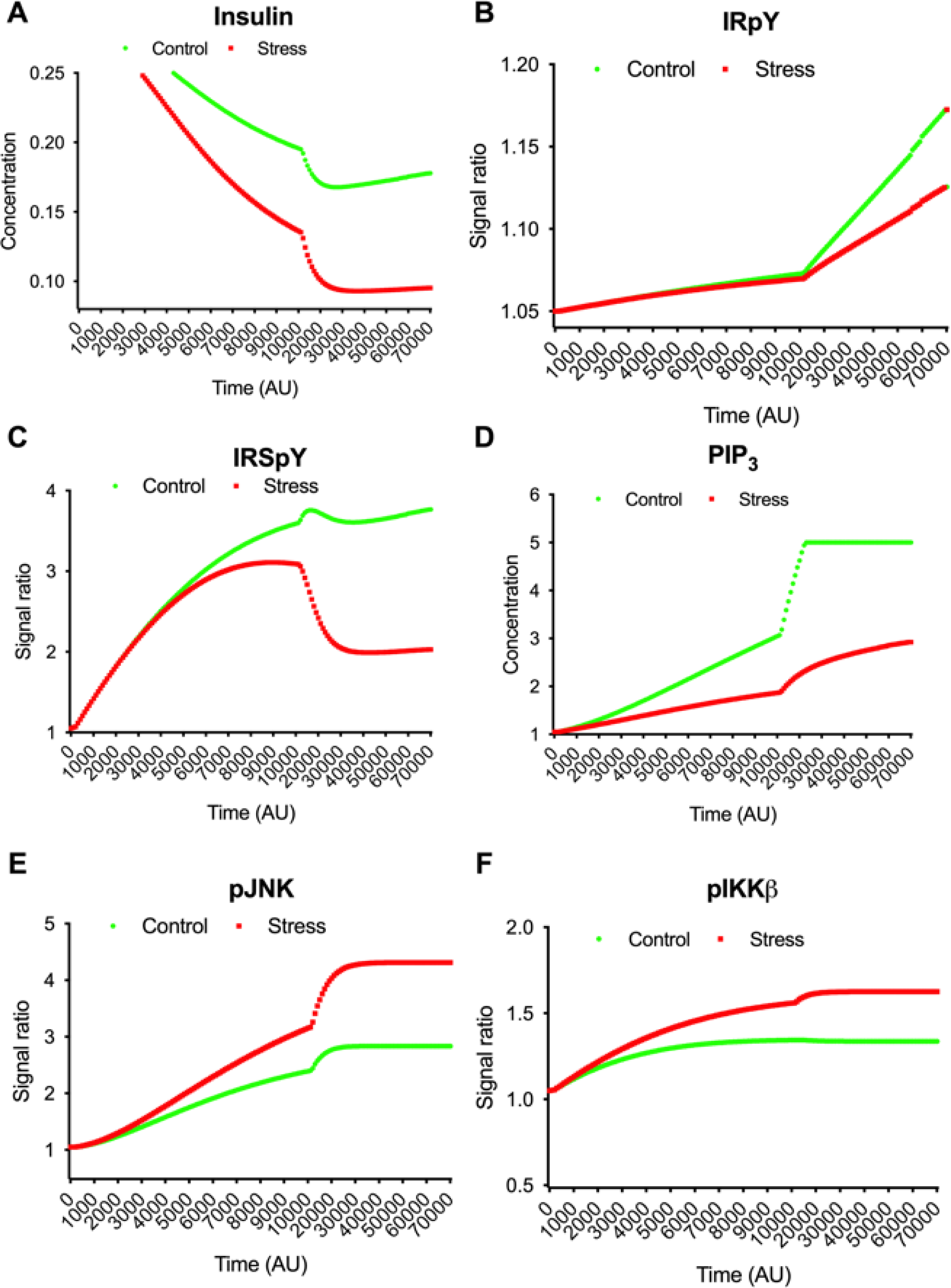
*In silico* state space model depicting that (A) insulin sensitivity is higher in control (NCD) than that in stress (DIO). Besides, signals of phosphorylated forms of intermediate molecules, (B) IRpY, (C) IRSpY and (D) concentration of PIP_3_ enhance in control (NCD) in comparison with stress (DIO) condition. On the other hand, the signals of (E) pJNK and (F) pIKKβ become higher during stress (DIO) than in control (NCD) scenario.

### Application of PID controller modeling CST activity

We observed that the state space model successfully mimicked the cellular behavior during both DIO and NCD conditions in consistent with the experimental results and existing literature. In addition, the aforesaid experimental results confirmed that CST alleviated ER stress. Literature showed ER stress would contribute to insulin resistance. Thus it indirectly implies that attenuation of ER stress in hepatic macrophages by CST may be an additional mechanism to enhance insulin sensitivity. In order to test the hypothesis, we applied two PID controllers on the state space model targeting pPERK, IRpY, IRSpY and pAKT to a high or low value so that relation between ER stress and insulin sensitivity on the application of CST could be depicted. We considered following three cases targeting i) pPERK and IRpY, ii) pPERK and IRSpY and iii) pPERK and pAKT.

#### Case 1 (Targeting pPERK and IRpY)

Here we checked all possible four conditions- i) High pPERK and low IRpY (DIO), ii) High pPERK and high IRpY (DIO), iii) Low pPERK and high IRpY (DIO+CST) and (iv) low pPERK and high IRpY (DIO+CST).

i. **High pPERK and low IRpY (DIO)** Here we set a high signal value for pPERK and a low signal value for IRpY (Fig. 7A-B). In this context, experimental results demonstrated that ER stress increased pPERK (Fig. 3B), peIF2α (Fig. 3C) and pIRE1α (Fig. 2B). In addition, the state space model showed decreased insulin sensitivity (Fig. 6A) and IRpY signal (Fig. 6B) in ER stress. Consistent with these results, this condition depicted high ER stress and low insulin sensitivity along with high values for the ratios, *i.e.*, (pPERK/PERK), (pIRE1α/IRE1α) and (peIF2α/eIF2α) (Fig. 7C-D). Thus, it depicts DIO condition.
ii. **High pPERK and high IRpY (DIO)** In this condition, both signals of pPERK and IRpY were set to high values (Fig. 7E-F). We observed that high signal of IRpY was not able to raise insulin sensitivity as ER stress became high (Fig. 7G). Subsequently, the other ratios (pPERK/PERK), (pIRE1α/IRE1α) and (peIF2α/eIF2α) became high (Fig. 7H). It is clear that when pPERK is high, *i.e.*, the ratios are high, insulin sensitivity cannot be high in spite of high IRpY signal. In consistent with the experimental as well as previous computational results, this situation also implies DIO condition.
iii. **Low pPERK and high IRpY (DIO+CST)** For this condition, the signal value of pPERK was targeted to be low while IRpY to be high (Fig. 8A-B). In this context, experimental results demonstrated CST alleviated ER stress (Fig. 4A-C and 5A-G), which led to decreased pPERK (Fig. 4G & I), peIF2α (Fig. 4G & J) and pIRE1α (Fig. 4G & H). Consistent with experimental results, this case depicted low ER stress (Fig. 8C) along with low values for the ratios (pPERK/PERK), (pIRE1α/IRE1α) and (peIF2α/eIF2α) (Fig. 8D). Thus this situation can be considered as DIO+CST condition. In addition, we noticed high insulin sensitivity (Fig. 8C) here.
iv. **Low pPERK and low IRpY (DIO+CST)** Here, we set low signal values for both pPERK and IRpY (Fig. 8E-F). In accordance with experimental results (Fig. 4–5), we observed very low ER stress (Fig. 8G) compared to DIO situation along with low values for the ratios (pPERK/PERK), (pIRE1α/IRE1α) and (peIF2α/eIF2α) (Fig. 8H). Thus this condition can also be considered as DIO+CST situation. Here we also observed that insulin sensitivity had been increasing slowly because of low IRpY signal (Fig. 8G).

**Figure 7:**
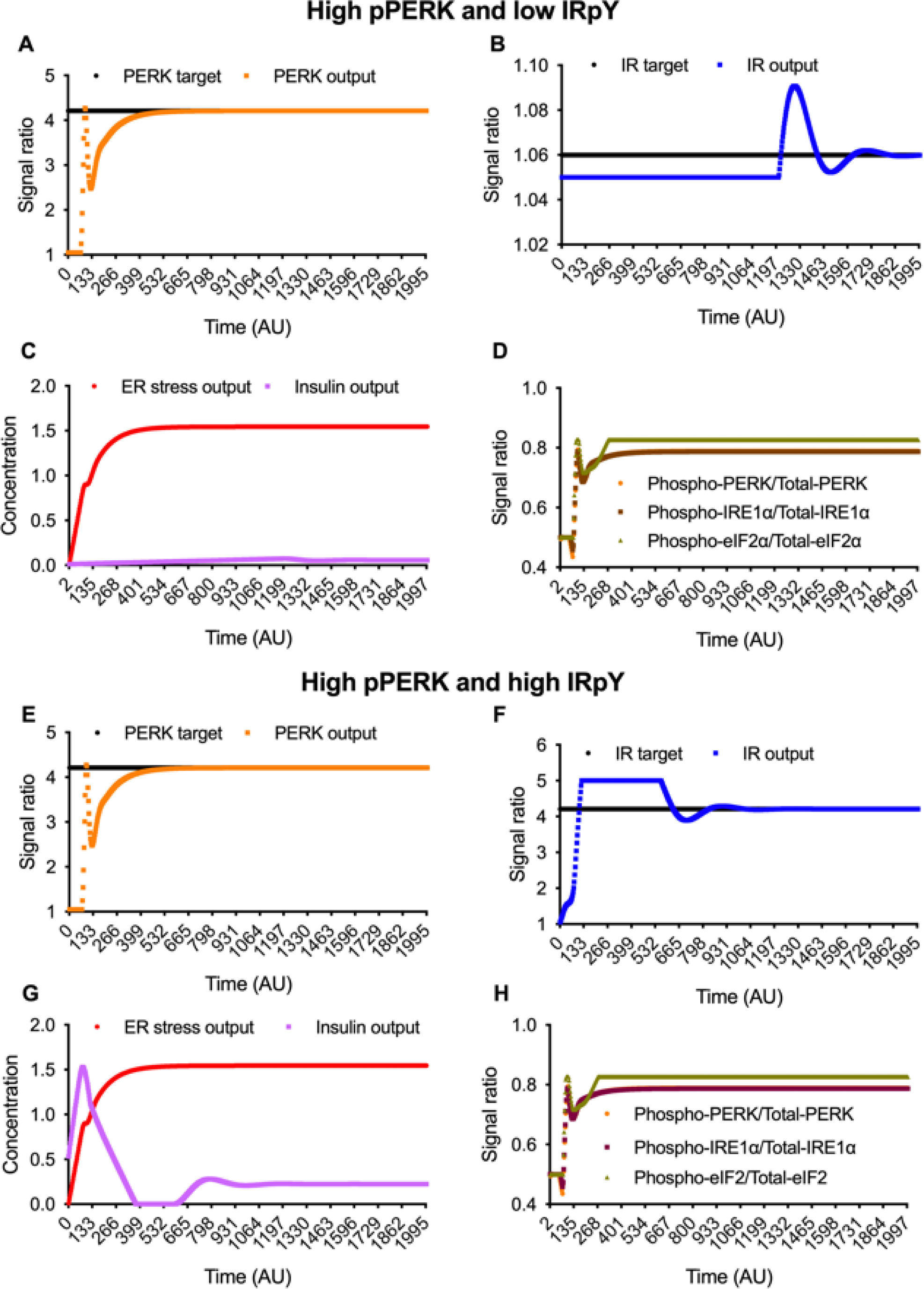
**High pPERK and low IRpY (DIO).** Here (A) phosphorylated PERK output as well as (B) tyrosine phosphorylated IR output is controlled by the PID controller according to the reference input PERK target and IR target respectively. As a result, (C) shows high ER stress and low insulin sensitivity. Besides, (D) ratios (phosphorylated PERK/total-PERK), (phosphorylatedIRE1α/total IRE1α) and (phosphorylated elF2α/total-elF2α) are quite high around 0.8. **High pPERK and high IRpY (DIO).** Here (E) phosphorylated PERK output as well as (F) tyrosine phosphorylated IR output is controlled by PID controller according to the reference input PERK target and IR target respectively. As a result, (G) shows high ER stress and low insulin sensitivity. Besides, (H) ratios (phosphorylated PERK/total PERK), (phosphorylated IRE1α/total IRE1α) and (phosphorylated eIF2α /total eIF2α) are quite high around 0.8.

**Figure 8:**
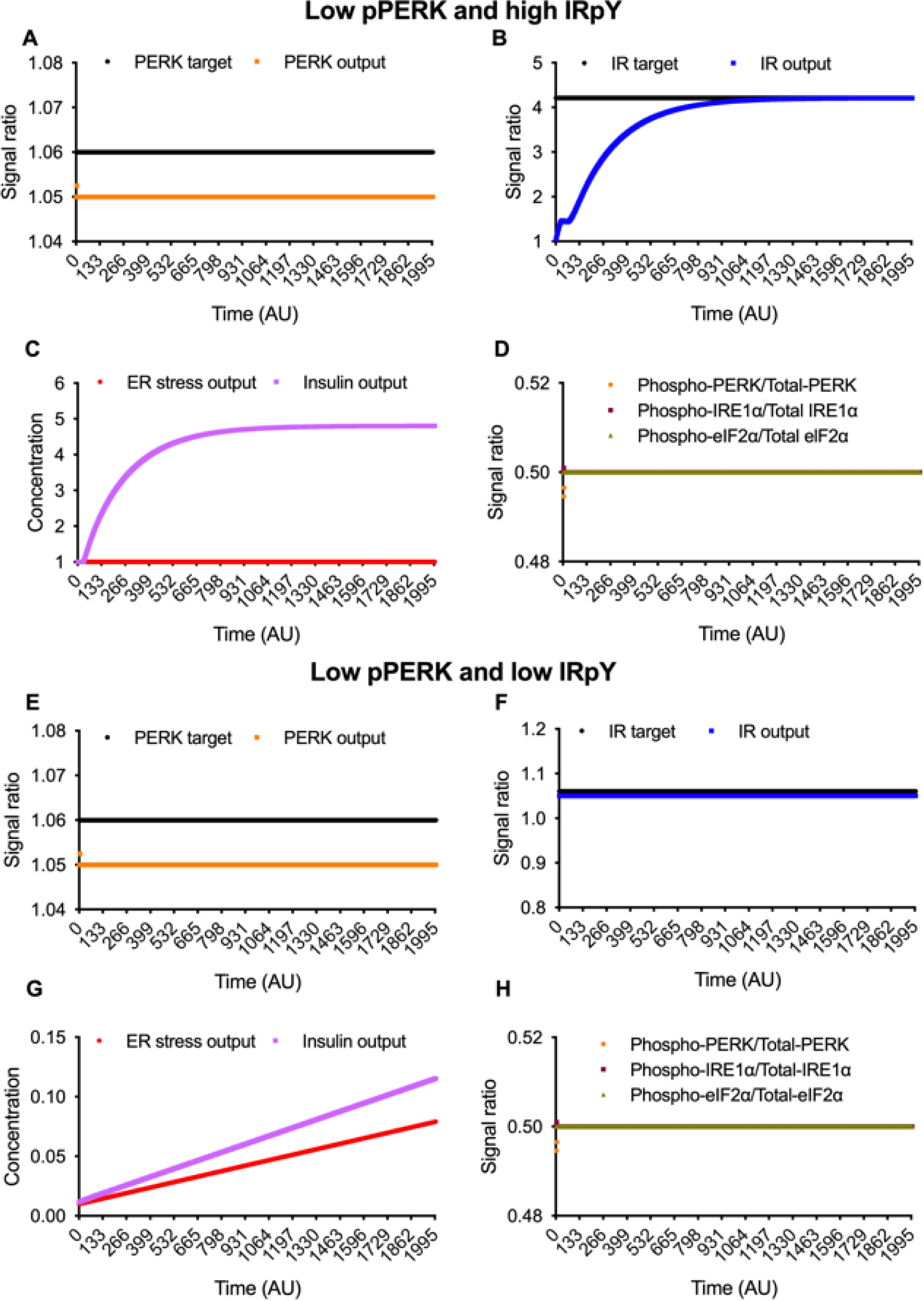
**Low pPERK and high IRpY (DIO+CST).** Here (A) phosphorylated PERK output as well as (B) tyrosine phosphorylated IR output is controlled by the PID controller according to the reference input PERK target and IR target respectively. As a result, (C) shows low ER stress and high insulin sensitivity. Besides, (D) ratios (phosphorylated PERK/total PERK), (phosphorylated IRE1α/total IRE1α) and (phosphorylated eIF2α/total eIF2α) are quite low around 0.5. **Low pPERK and low IRpY (DIO+CST).** Here (E) phosphorylated PERK output as well as (F) tyrosine phosphorylated IR output is controlled by the PID controller according to the reference input PERK target and IR target respectively. As a result, (G) shows that ER stress is very low compared to that under DIO condition, while insulin sensitivity is increasing. Besides, (H) ratios (phosphorylated PERK/total PERK), (phosphorylated IRE1α/total IRE1α) and (phosphorylated eIF2α/Total eIF2α) are quite low around 0.5.

From the above four conditions, we clearly observed enhanced insulin sensitivity in DIO+CST scenario. Besides, low ER stress and low values of the ratios (pPERK/PERK), (pIRE1α/IRE1α) and (peIF2α/eIF2α) would result in high insulin sensitivity. Thus we can conclude that CST not only reduces ER stress but also increases insulin sensitivity. Higher level of IRpY could not enhance insulin sensitivity when ER stress is high.

#### Case 2 (Targeting pPERK and IRSpY)

Similar to case 1, we checked all possible four conditions here- i) High pPERK and low IRSpY (DIO), ii) High pPERK and high IRSpY (DIO), iii) Low pPERK and high IRSpY (DIO+CST) and (iv) low pPERK and high IRSpY (DIO+CST).

i. **High pPERK and low IRSpY (DIO)** Here a high signal value for pPERK and a low signal value for IRSpY (Fig. S4A-B of Supplementary File S1) led to high ER stress and low insulin sensitivity along with high values for the ratios, *i.e.*, (pPERK/PERK), (pIRE1α/IRE1α) and (peIF2α/eIF2α) (Fig. S4C-D of Supplementary File S1). Thus, it depicts DIO condition.
ii. **High pPERK and high IRSpY (DIO)** High signal of IRSpY (Fig. S4F of Supplementary File S1) was not able to raise insulin sensitivity (Fig. S4G of Supplementary File S1) because of high pPERK (Fig. S4E of Supplementary File S1) resulting in high ER stress (Fig. S4G of Supplementary File S1). Subsequently, the other ratios (pPERK/PERK), (pIRE1α/IRE1α) and (peIF2α/eIF2α) became high (Fig. S4H of Supplementary File S1). Similarly, it also implies DIO condition.
iii. **Low pPERK and high IRSpY (DIO+CST)** When the signal value of pPERK was targeted to be low while IRSpY to be high (Fig. S5A-B of Supplementary File S1), we noticed high insulin sensitivity (Fig. S5C of Supplementary File S1). Besides, it depicted low ER stress (Fig. S5C of Supplementary File S1) along with low values for the ratios (pPERK/PERK), (pIRE1α/IRE1α) and (peIF2α/eIF2α) (Fig. S5D of Supplementary File S1). This situation arises when CST is applied.
iv. **Low pPERK and low IRSpY (DIO+CST)** Similarly, low signal value for pPERK (Fig. S5E of Supplementary File S1) led to increasing insulin sensitivity (Fig. S5G of Supplementary File S1). However, insulin sensitivity had been increasing slowly because of low IRSpY signal (Fig. S5F of Supplementary File S1). Besides, we observed very low ER stress (Fig. S5G of Supplementary File S1) compared to DIO situation along with low values for the ratios (pPERK/PERK), (pIRE1α/IRE1α) and (peIF2α/eIF2α) (Fig. S5H of Supplementary File S1). This condition also represents DIO+CST scenario.

Thus, reduction of pPERK on application of CST is the main factor to enhance insulin sensitivity. Here, signal level of IRSpY has no role to enhance insulin sensitivity when ER stress is high.

#### Case 3 (Targeting pPERK and pAKT)

Similar to previous two cases, we investigated all possible four conditions- i) High pPERK and low pAKT (DIO), ii) High pPERK and high pAKT (DIO), iii) Low pPERK and high pAKT (DIO+CST) and (iv) Low pPERK and high pAKT (DIO+CST).

i. **High pPERK and low pAKT (DIO)** We found high ER stress and low insulin sensitivity along with high values for the ratios, *i.e.*, (pPERK/PERK), (pIRE1α/IRE1α) and (peIF2α/eIF2α) (Fig. S6C-D of Supplementary File S1) due to a high signal value for pPERK and a low signal value for pAKT (Fig. S6A-B of Supplementary File S1). As in the previous two cases, it depicts DIO condition.
ii. **High pPERK and high pAKT (DIO)** High signal of pAKT (Fig. S6F of Supplementary File S1) was able to enhance insulin sensitivity (Fig. S6G of Supplementary File S1) in spite of high pPERK (Fig. S6E of Supplementary File S1) resulting in high ER stress (Fig. S6G of Supplementary File S1). Besides, the other ratios (pPERK/PERK), (pIRE1α/IRE1α) and (peIF2α/eIF2α) remained high (Fig. S6H of Supplementary File S1). It is also a DIO condition.
iii. **Low pPERK and high pAKT (DIO+CST)** Low signal value of pPERK and high signal value of pAKT (Fig. S7A-B of Supplementary File S1), enhanced insulin sensitivity (Fig. S7C of Supplementary File S1). Besides, ER stress became low (Fig. S7C of Supplementary File S1). This condition also depicted low values for the ratios (pPERK/PERK), (pIRE1α/IRE1α) and (peIF2α/eIF2α) (Fig. S7D of Supplementary File S1). Experimental results showed that CST reduced both pPERK and ER stress.
iv. **Low pPERK and low pAKT (DIO+CST)** Here, low signal value for pPERK (Fig. S7E of Supplementary File S1) led to improved insulin sensitivity (Fig. S7G of Supplementary File S1) in spite of low pAKT signal (Fig. S7F of Supplementary File S1). Besides, very low ER stress (Fig. S7G of Supplementary File S1) compared to DIO situation along with low values for the ratios (pPERK/PERK), (pIRE1α/IRE1α) and (peIF2α/eIF2α) (Fig. S7H of Supplementary File S1) were found. Obviously, this condition can be considered as DIO+CST condition.

Finally, it can be concluded that although reduction of pPERK on application of CST drives enhanced insulin sensitivity, pAKT can be treated as another drug target during high ER stress. High signal level of pAKT may enhance insulin sensitivity in spite of high ER stress.

## Discussion

Experimental analyses revealed that chronic treatment of DIO mice with CST results in attenuation of ER stress. How does CST alleviate ER stress? Inflammation has been reported to influence ER stress (*88*, *89*). Conversely, ER stress has been implicated in several inflammatory diseases including obesity and diabetes, inflammatory bowel diseases, arthritis and spondyloarthropathies, neurodegenerative and neuromuscular inflammatory diseases, and numerous forms of respiratory inflammation (*65*, *90*–*94*). UPR signaling and ER stress utilize the following mechanisms to influence inflammation: (i) Protein misfolding triggers the translocation of ATF6α from the ER to the Golgi destined for its cleavage by the S1P and S2P proteases (*38*). Consequently, increased phosphorylation of AKT activates NF-κB through the release and translocation of the active transcription factors to the nucleus (*95*); (ii) Autophosphorylated PERK-induced phosphorylation of eIF2α inhibits protein translation and reduce IκBα production, and thereby, influence transcription of NF-κB (*96*, *97*), and (iii) Upon binding to the adaptor protein TNF-receptor activating factor 2 (TRAF2) phosphorylated IRE1α activates the JNK/AKT pathway and phosphorylate IKKβ, which leads to cleavage of Ikβα and activation of NF-κB (*98*). Our findings revealed that treatment of DIO mice with CST resulted in marked decrease in phosphorylation of PERK, eIF2α, IRE1α and NF-κB. Therefore, one of the mechanisms by which CST can alleviate ER stress is by reducing inflammation as shown here as well as shown previously in DIO liver (*31*), in macrophage-driven atherosclerosis (*99*), in two mouse models of experimental colitis (*100*) as well as in reactivation of intestinal inflammation in adult mice with previously established chronic colitis (*101*).

ER stress could contribute to the development of insulin resistance utilizing the following pathways: (i) UPR activated transcription factors modulate expression of the gluconeogenic enzymes PEPCK and G6Pase as well as lipogenic enzyme SREBP1c. It has been reported that ER stress increases glucose-6-phosphatase activity and glucose output in primary hepatocytes (*102*). This decreased insulin signaling was mediated by activated IRE1, probably through TRAF2 recruitment and JNK activation (*84*). Furthermore, gluconeogenesis could be activated either by ER stress-induced inhibition of IL-6/STAT3-dependent suppression of hepatic gluconeogenic enzyme expression (*103*) or by activated (by ER stress) CREBH-induced augmentation of transcription of gluconeogenic genes (*104*). We have reported recently that CST decreased gluconeogenesis by inhibiting expression of PEPCK and G6Pase genes (*31*). (ii) ER stress-induced activation of IRE1 recruits JNK and IKK by recruiting TRAF2 and ASK1 (*84*, *87*), which impair insulin signaling by phosphorylating IRS1 on serine residues. Furthermore, saturated fatty acids, ceramides and ER stress activate PKR, which inhibits insulin signaling by inducing phosphorylation of serine residues in IRS1 (direct regulation) as well as activating JNK signalling pathway (indirect regulation) (*63*, *105*). Finally, the PERK-induced activation of TRB3, the ER stress-inducible tribbles ortholog in humans also leads to the impairment of insulin signaling by inhibiting AKT (PKB) (*106*). Furthermore, NF-κB can also be activated by ER stress-induced PERK pathway and ATF6 branches (*95*). (iii) UPR promotes accumulation of fat in hepatocytes by inducing *de novo* lipogenesis (direct effect) and affecting VLDL secretion (indirect effect) resulting in the development of insulin resistance. Here, we found that CST decreased ER lumen diameter in DIO liver macrophages, indicating decreased ER stress. Thus, attenuation of ER stress in hepatic macrophages by CST may also contribute to improved insulin sensitivity in addition to inhibition of Ly6C+ macrophage infiltration and inflammation as well as suppression of hepatic glucose production (*31*).

To test the above hypothesis, we developed a PID controller based state space model. Firstly, an *in silico* state space model was designed by integrating ER stress and insulin signaling pathway, and validated with the experimental results for both DIO and NCD conditions. The present state space model resembled the experimental behavioral pattern including high ratios of (pPERK/PERK), (pIRE1α/IRE1α) and (peIF2α/eIF2α) during ER stress (DIO). In addition, in consistent with existing literature, the state space model showed decreased insulin sensitivity along with decreased IRpY, IRSpY, pAKT and pFoxO1 signals along with concentration of PIP_3_ due to the activation of pJNK and pIKKβ during ER stress (DIO). Secondly, we applied two PID controllers on the present state space model to draw a relation between ER stress and insulin sensitivity during DIO+CST situation. PID controller based simulation followed the experimentally observed behavior of ER stress along with the ratios, (pPERK/PERK), (pIRE1α/IRE1α) and (peIF2α/eIF2α), during both DIO and DIO+CST conditions. Subsequently, it also showed that insulin sensitivity would enhance during DIO+CST scenario. Even higher levels of tyrosine phosphorylations of IR and IRS were not able to raise insulin sensitivity during high ER stress. On the other hand, according to *in silico* studies, high level of phosphorylated AKT enhanced insulin sensitivity in spite of high ER stress. This prediction generated by our *in silico* model awaits experimental verification. However, on application of CST, reduced pPERK, resulting in low ER stress, can increase insulin sensitivity independent of pAKT signal. Thus, we can conclude that CST reduces not only inflammation but also the ER stress and, as a result, increases the insulin sensitivity in obese model. Additionally, we can treat AKT as another drug target to enhance insulin sensitivity during high ER stress.

## Acknowledgements

The electron micrographs were taken in the Cellular and Molecular Medicine Electron microscopy core facility at UCSD, which is supported in part by National Institute of Health Award number S10OD023527. Abhijit Dasgupta, one of the authors, gratefully acknowledges Ministry of Electronics and Information Technology, Government of India, for providing him a senior research fellowship under Visvesvaraya PhD scheme for Electronics and IT. A part of this work was done, when one of the authors, Rajat K. De, was visiting University of California, San Diego, USA, under Fulbright-Nehru Academic and Professional Excellence Fellowship Program. This research was supported mainly by S.K.M.’s personal fund and partly by a grant from the Department of Veterans Affairs (I01BX000323 to S.K.M.).

## Author contributions statement

AD and RKD conceived the idea of modeling the integrated insulin signaling and ER stress pathway to investigate CST effects on ER stress and insulin sensitivity. IR provided some biological input to the model. SKM conceived the idea of CST alleviation of ER stress, performed *in vivo* experiments and made the cartoons and graphics. KB performed *in vivo* experiments. GKB helped in analyzing results.

## Additional Information

There is NO Competing Interest.

